# Parkinsonian rest tremor can be distinguished from voluntary hand movements based on subthalamic and cortical activity using machine learning

**DOI:** 10.1101/2023.02.07.527275

**Authors:** Dmitrii Todorov, Alfons Schnitzler, Jan Hirschmann

## Abstract

Tremor is one of the cardinal symptoms of Parkinson’s disease. The neurophysiology of tremor is not completely understood, and so far it has not been possible to distinguish tremor from voluntary hand movements based on local brain signals.

Here, we re-analyzed magnetoencephalography and local field potential recordings from the subthalamic nucleus of six patients with Parkinson’s disease. Data were obtained after withdrawal from dopaminergic medication (Med Off) and after administration of levodopa (Med On). Using gradient-boosted tree learning, we classified epochs as tremor, self-paced fist-clenching, static forearm extension or tremor-free rest.

While decoding performance was low when using subthalamic activity as the only feature (balanced accuracy mean: 38%, std: 7%), we could distinguish the four different motor states when considering cortical and subthalamic features (balanced accuracy mean: 75%, std: 17%). Adding a single cortical area improved classification by 17% on average, as compared to classification based on subthalamic activity alone. In most patients, the most informative cortical areas were sensorimotor cortical regions. Decoding performance was similar in Med On and Med Off.

Our results demonstrate the advantage of monitoring cortical signals in addition to subthalamic activity for movement classification. By combining cortical recordings, subcortical recordings and machine learning, future adaptive systems might be able to detect tremor specifically and distinguish between several motor states.

## Introduction

Deep brain stimulation (DBS) is a widely used treatment for advanced Parkinson’s disease (PD; Krauss et al., 2021). While well accepted and effective, it also has significant side-effects (Zarzycki and Domitrz, 2020) most of which result from current spread to structures adjacent to the DBS target (Koeglsperger et al., 2019). Hence, a general strategy for reducing side-effects is to reduce the energy applied in DBS. One way to achieve this while maintaining clinical benefits is to adapt stimulation to the current motor state. This approach is called adaptive DBS (Meidahl et al., 2017; Neumann et al., 2019; Krauss et al., 2021). Various signals can be used to control either the onset or the amplitude of DBS (Marceglia et al., 2021). Currently, the most investigated neural control signal is subthalamic beta band activity (Little et al., 2013, 2016; Piña-Fuentes et al., 2017; Tinkhauser et al., 2017), but other signals have also been considered, such as cortical gamma band activity (Swann et al., 2018; Gilron et al., 2021) or local evoked potentials (Dale et al., 2022).

Adapting DBS to the current motor state makes sense for treating tremor, particularly, because tremor is highly variable, waxing and waning spontaneously. In case of tremor, one natural strategy for adaptive DBS is to monitor tremor directly by means of peripherals such as accelerometers or electromyography (Malekmohammadi et al., 2016a; Cagnan et al., 2017; Cernera et al., 2021). This approach, however, requires additional hardware and thus raises additional security and compliance issues.

An alternative strategy is to use brain signals rather than peripherals. It is known that Parkinsonian rest tremor originates in the brain (Milosevic et al., 2018), involving an extended subcortico-cortical network including the basal ganglia, cerebellum, thalamus and motor cortex (Timmermann et al., 2003; Helmich, 2018). Thus, it should be feasible to track tremor by monitoring its neural correlates. Several studies have investigated subcortical correlates by analyzing local field potentials (LFPs) recorded from DBS electrodes and identified numerous tremor-related changes, such as an increase of power at individual tremor frequency (Hirschmann et al., 2013a), a beta power decrease (Wang et al., 2005; Qasim et al., 2016), an increase of low gamma power (Beudel et al., 2015), and a shift of power in the very high frequency range (Hirschmann et al., 2016). While these studies were able to distinguish tremor from tremor-free rest periods, no study has been able to distinguish PD rest tremor from voluntary hand movements so far. When applying adaptive DBS in practice, it would be desirable to make this distinction to reduce unnecessary stimulation.

In this study, we re-analyzed simultaneous recordings of forearm EMG, magnetoencephalography (MEG) and subthalamic nucleus LFP data, obtained from tremor-dominant PD patients, aiming to distinguish between PD rest tremor, self-paced fist-clenching, static forearm extension and tremor-free rest periods (referred to as “quiet” episodes from here on), by means of gradient-boosted tree learning, a popular method in machine learning. We show that these motor states can be distinguished when considering cortical and subthalamic data.

## Methods

### Participants

In this study we used a subset of previously collected data (Hirschmann et al., 2013a). We selected patients with enough data for comparing rest tremor, quiet episodes, and two motor tasks (see below). Recordings from 6 patients met these requirements. Overall, we had 11 datasets (5 subjects in Med On and Med Off, one subject in Med Off only; see Tab. 1).

**Tab. 1:** Patient data. In the “Electrode model” column, M: 4-contact, non-segmented electrode by Medtronic (model 3389), S: 4-contact, non-segmented electrode by St. Jude Medical.

| subj | Gender | Age (y) | Disease duration (y) | Tremor frequency (Hz) | Electrode model | Medication states recorded |
| --- | --- | --- | --- | --- | --- | --- |
| S01 | m | 65 | 8 | 4 | M | ON, OFF |
| S02 | m | 69 | 6 | 3.5 | M | ON, OFF |
| S03 | m | 68 | 11 | 3 | M | OFF |
| S04 | m | 68 | 2 | 4 | S | ON, OFF |
| S05 | m | 52 | 11 | 6 | S | ON, OFF |
| S07 | m | 59 | 6 | 4.5 | M | ON, OFF |

All patients involved were diagnosed with idiopathic PD, and underwent DBS surgery the day before measurement. They experienced spontaneous waxing and waning of rest tremor during the recordings. Patient details are provided in Tab. 1 The study was approved by the ethics committee of the Medical Faculty of the Heinrich Heine University Düsseldorf (Study No. 3209). It was carried out in accordance with the Declaration of Helsinki and required written informed consent.

### Recordings

Patients were recorded one day after STN electrode implantation with leads still externalized. All but one patients were recorded in two sessions: one session OFF oral dopaminergic medication for at least 12h and one session ON medication; one patient had only the OFF medication session recorded. Subcutaneous apomorphine administration was paused 1.5-2h before measurements started. Each session contained four parts: rest, motor task 1, rest, motor task 2. Rest periods lasted 5 min. Motor task 1 was static forearm extension (“hold”) and motor task 2 was self-paced fist-clenching (“grasp”) at approximately 1 Hz (Hirschmann et al., 2013b). Movements were performed with the symptom-dominant body side in five 1-min blocks which were interleaved by 1 min pauses to avoid fatigue. During rest and between the blocks of voluntary movement tremor appeared and disappeared spontaneously.

Local field potentials from the STN, the magnetencephalogram (MEG; Vectorview, MEGIN) and the surface electromyogram of the extensor digitorum communis and flexor digitorum superficialis muscles of both upper limbs were recorded simultaneously. The sampling rate was 2000 Hz. DBS electrodes were connected to the amplifier of the MEG system by externalized, non-ferromagnetic leads. Electrode contacts were referenced to the left mastoid and rearranged to a bipolar montage offline by subtracting signals from neighboring contacts. EMG electrodes were referenced to surface electrodes at the muscle tendons. A hardware filter was applied with a pass-band of 0.1–660 Hz.

### EMG labeling

The hand performing the voluntary movements was presenting intermittent rest tremor in all subjects. In order to label the data, EMG data was 10Hz high-pass filtered, rectified and smoothed with a 100ms box-filter. Quiet periods were identified semi-automatically. Epochs with both EMG channels (forearm flexor and extensor) deflecting less than their respective 40% quantiles (computed using the data from the entire recording) were labeled as candidate quiet periods automatically and then adapted manually. Tremor and voluntary movements were labeled manually. Epochs with uncertain labels, as well as postural and kinetic tremor were discarded. When possible, “safety offsets” were included to ensure that different behavioral states were separated by at least 1s to mitigate the risk of confusing states (Fig. 1). The amount of data available per motor state is provided in Tab. 2.

**Tab. 2.** Amount of data per motor state (after discarding MEG and LFP artifacts) in seconds and balanced accuracy (bacc) in % for the feature set consisting of Hjorth activity of the best LFP channel and all cortical areas. The last column identifies the hand that was used to define the behavioral states.

| medication OFF |  |  |  |  |  | medication ON |  |  |  |  |  |
| --- | --- | --- | --- | --- | --- | --- | --- | --- | --- | --- | --- |
| subject | tremor | rest | hold | grasp | bacc | tremor | rest | hold | grasp | bacc | hand |
| S01 | 240 | 384 | 307 | 238 | 91% | 13 | 584 | 282 | 219 | 81% | left |
| S02 | 413 | 423 | 186 | 310 | 95% | 350 | 169 | 311 | 320 | 79% | left |
| S03 | 16 | 434 | 335 | 253 | 45% | N/A | N/A | N/A | N/A | N/A | left |
| S04 | 56 | 403 | 252 | 119 | 73% | 54 | 426 | 279 | 94 | 64% | left |
| S05 | 353 | 510 | 121 | 324 | 89% | 18 | 539 | 293 | 261 | 87% | right |
| S07 | 77 | 451 | 258 | 236 | 47% | 427 | 328 | 284 | 286 | 83% | left |
| mean | 192.5 | 434.2 | 243.2 | 246.67 | 73% | 172.4 | 409.2 | 289.8 | 236 | 78% |  |
| std | 167.19 | 43.92 | 78.74 | 72.828 | 22% | 199.77 | 167.3 | 12.95 | 87.51 | 8% |  |

**Figure 1.**
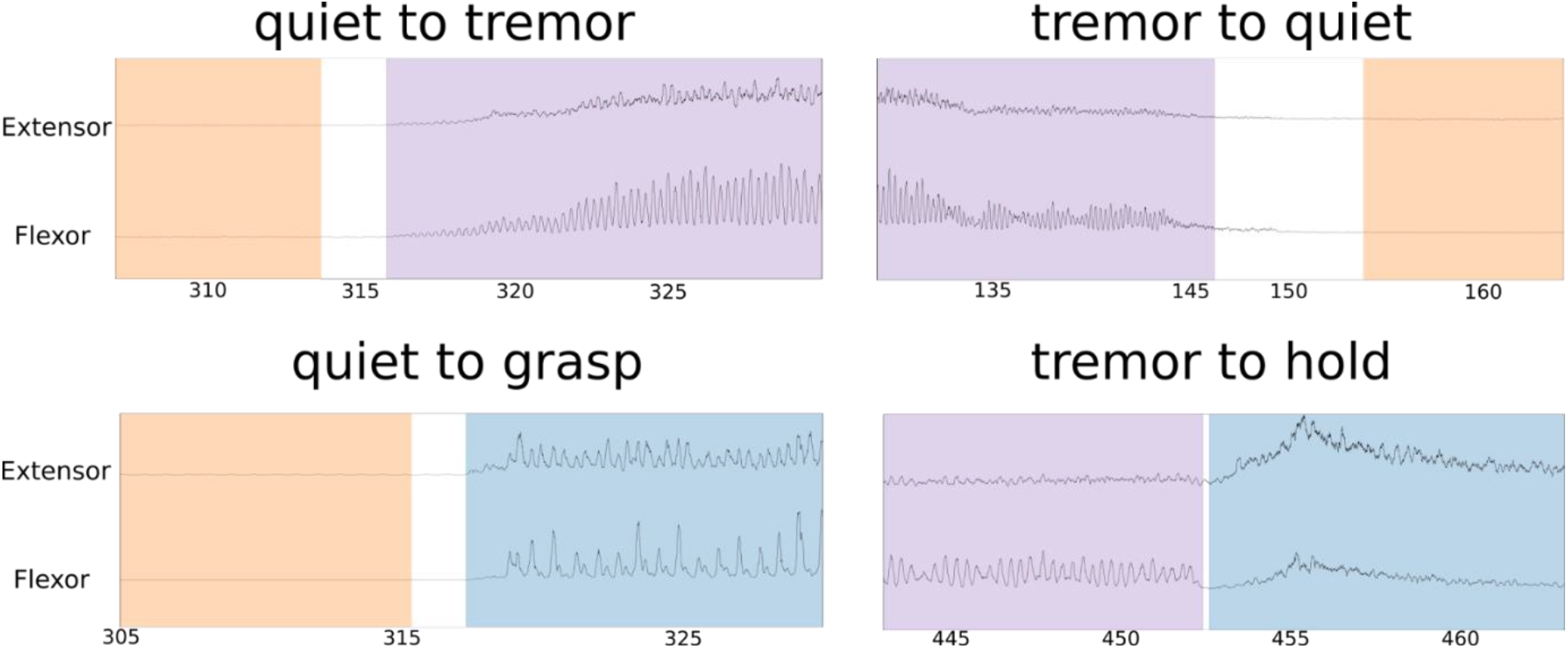
Labeling of behavioral states in forearm EMG, state transition examples.

### Preprocessing

The MEG sensor data and LFP data were resampled to 256Hz to increase computation speed. MEG artifacts were identified as time periods when sensor data deflected more than 2.5 times the signal’s 95%-trimmed mean in multiple sensors. Spatiotemporal signal space separation (Taulu and Simola et al., 2006) with a10s time window was applied to the MEG data, using the MNE toolbox (Gramfort et al. 2014). LFP artifacts were identified as periods with strong broadband modulation of the signal.

### Source reconstruction

For forward modelling, we made use of realistic, single-shell head models based on the individual, T1-weighted MR image (Nolte, 2003). Source reconstruction was performed by means of Linearly Constrained Minimum Variance (LCMV) beamforming, as implemented in the Fieldtrip toolbox (Oostenveld et al., 2011). The beamformer grid contained 567 locations, spread out evenly across the cortical and cerebellar surface. It was aligned to Montreal Neurological Institute (MNI) space, allowing for grid parcellation into 30 supersets of regions defined in the Automatic Anatomic Labeling (AAL) atlas (Tzourio-Mazoyer et al., 2002). For the computation of the spatial filter, we used only the quiet periods without artifacts. For LCMV regularization, the lambda parameter was set to 0.1% of the maximum eigenvalue of the covariance matrix computed on the quiet periods.

### Feature engineering

For feature engineering, the data were separated into 1s-long disjoint windows. Windows containing MEG or LFP artifacts were discarded. The number of windows per dataset averaged across subjects and medication conditions after artifact and uncertain EMG state rejection was 1112 (std = 172), whereas total recording duration (without artifact rejection) was 1638s (std = 168s). For each window, we computed the logarithm of Hjorth activity, equal to windowed signal variance <u>(Hjorth 1970)</u>. The logarithm served to bring the data distribution closer to normal. For the source-reconstructed data, we averaged Hjorth activity across all grid points belonging to the same cortical parcel to obtain one feature per cortical area and time window.

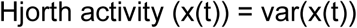

The use of Hjorth activity rather than spectral power or connectivity was motivated by the intention to compare decoding performance across brain areas while keeping the number of features as low as possible to avoid overfitting. Thus, we preferred metrics without frequency-resolution, uniquely associated with a single area. Our analysis choices were guided by the work of Yao et al. who performed a detailed investigation of the optimal choices for automated tremor detection (Yao et al., 2020). The authors tested several features and identified Hjorth activity as one of the most useful features. Further, they found XGBoost to be the best performing machine learning model among several candidates, and 1s to be a good window size.

### Machine Learning

To predict the four different motor states from Hjorth activity, we performed 4-label classification on the windowed data using 5-fold, shuffled cross-validation. The XGBoost algorithm was used to perform the classification. XGBoost is an advanced decision-tree-based algorithm, widely used machine learning (Chen and Guestrin, 2016). It often provides better results than linear models and requires relatively small amounts of training data. Since the number of windows differed substantially between behavioral states both within and between subjects (Tab. 2), we used balanced accuracy, defined as the mean of sensitivity and specificity averaged across classes, as a measure of performance (Kelleher et al., 2015). When training the model, we used oversampling via *imbalanced-learn* toolbox (Lemaître et al., 2017) to artificially balance classes. The entire python & Matlab pipeline code (including preprocessing, source reconstruction and machine learning) is available at the code repository github.com/todorovdi/ContBehFeatExplorer.

### LFP channel selection and hyperparameter tuning

For each subject, we performed single-feature classification for each LFP channel separately and selected the channel with best out-of-sample performance. Similarly, hyperparameter tuning was done by sampling combinations of the XGboost parameters *max_depth, min_child_weight, subsample* and *eta* and selecting the combination giving the best 5-fold cross-validated balanced accuracy. Neither LFP channel selection nor tuning had a strong effect on performance. All selected LFP channels were located contralateral to movement, except for patient S05.

## Results

### Distinguishing movements based on STN activity

We investigated how well the different motor states could be distinguished based on subthalamic activity alone. Fig. 2 shows a confusion matrix for each subject and medication state, providing discrimination performance for each pair of movements. STN activity alone turned out to be a rather weak predictor of movement type. In medication OFF (N=6), the average percentage of correctly classified windows was 62% for quiet (std=5%), 32% for rest tremor (std=19%), 34% for hold (std=16%), and 23% for grasp (std=27%). The most common misclassification was labeling tremor as hold (mean number of cases=22%, std=9%). The least common misclassification was labeling tremor as grasp (mean=8%, std=8%). Random shuffling of labels lowered performance, demonstrating that the STN did provide relevant information (Tab. 3).

**Figure 2:**
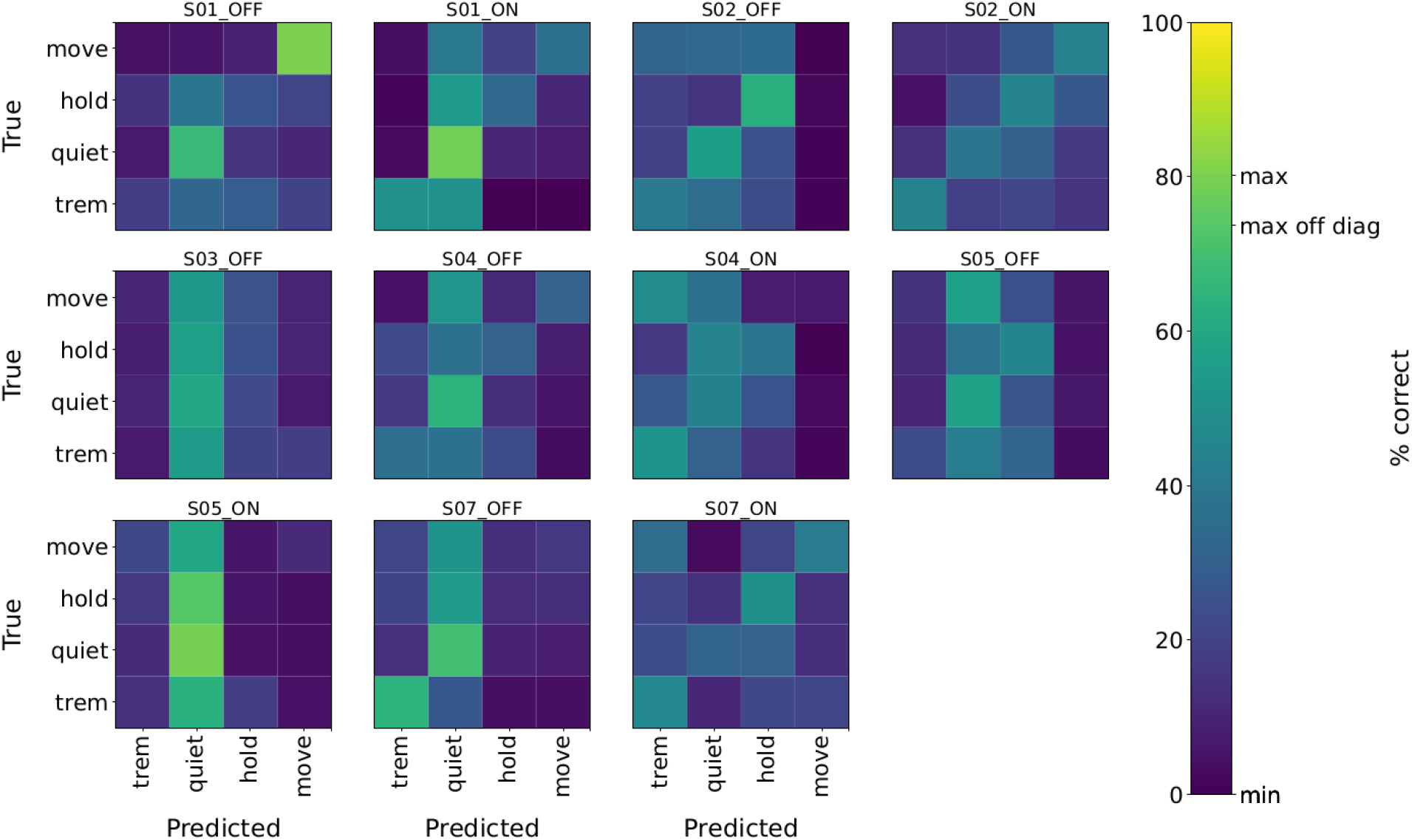
Distinguishing movements based on subthalamic activity. Confusion matrices for Med Off and On per subject, averaged across cross-validation folds. The color bar shows the overall maximum and minimum, the maximum of all off-diagonal entries and the minimum of all diagonal entries.

**Tab. 3.** Balanced accuracy, group results. “Shuffled” performance is 90% percentile of balanced accuracies obtained from 100 permutations of class labels.

| Data used: STN LFP alone |  |  |  |  | Data used: STN LFP + MEG |  |  |  |  |
| --- | --- | --- | --- | --- | --- | --- | --- | --- | --- |
| medication | classification | shuffling | mean, % | std, % | medication | classification | shuffling | mean, % | std, % |
| OFF | all vs. all | no | 38 | 8 | OFF | all vs. all | no | 73 | 22 |
| OFF | all vs. all | yes | 26 | 1 | OFF | all vs. all | yes | 25 | 1 |
| OFF | trem vs. all other | no | 59 | 8 | OFF | trem vs. all other | no | 78 | 19 |
| OFF | trem vs. all other | yes | 50 | 0 | OFF | trem vs. all other | yes | 50 | 0 |
| OFF | trem vs. rest | no | 64 | 4 | OFF | trem vs. rest | no | 80 | 16 |
| OFF | trem vs. rest | yes | 51 | 1 | OFF | trem vs. rest | yes | 50 | 0 |
| ON | all vs. all | no | 40 | 8 | ON | all vs. all | no | 79 | 9 |
| ON | all vs. all | yes | 26 | 0 | ON | all vs. all | yes | 25 | 1 |
| ON | trem vs. all other | no | 59 | 6 | ON | trem vs. all other | no | 86 | 8 |
| ON | trem vs. all other | yes | 50 | 0 | ON | trem vs. all other | yes | 50 | 0 |
| ON | trem vs. rest | no | 60 | 8 | ON | trem vs. rest | no | 87 | 8 |
| ON | trem vs. rest | yes | 51 | 1 | ON | trem vs. rest | yes | 51 | 1 |

Levodopa did not have a strong influence on movement discrimination (Tab. 3). Balanced accuracy did not differ significantly between medication states (*p* = 0.7, all vs. all, independent t-test, NOFF = 6, NON =5).

In addition to the multiclass problem, we tested how well our approach can discriminate tremor from any other movement, as this classification is relevant for tremor therapy through adaptive DBS. When merging quiet, hold and grasp epochs into a single non-tremor class, we obtained balanced accuracy (in this case of two classes it is equal to the mean of sensitivity and specificity) of 59% in the Med Off state (std=8%) and 58% in Med On (std=6%; Tab. 2). When discriminating merely between rest tremor and quiet, accuracy was 64% in Med Off (std=4%) and 60% in Med On (std=7%). This performance is similar to previous papers dealing with tremor vs. rest (Hirschmann et al., 2017; Yao et al., 2020), taking into account that they reported regular (not balanced) accuracy.

### Distinguishing movements based on STN and cortical activity

When classifying epochs based on both subthalamic activity and the activity of all cortical parcels, we observed a strong increase in performance (average performance gain: 37%; Tab. 3 and Fig. 3). In medication OFF (N=6), the average percentage of correctly classified epochs was 74% for quiet (std=21%), 58% for rest tremor (std=36%), 80% for hold (std=12%), and 81% for grasp (std=18%). The most common misclassification was labeling tremor as hold (mean=22%, std=31%). The least common misclassification was labeling rest tremor as grasp (mean=1%, std=1%). As in the STN-only case, random shuffling of labels lowered performance, lowering the number of classes increased performance, and medication had little influence on performance, independent t-test p=0.63 (see also Tab. 3).

**Figure 3:**
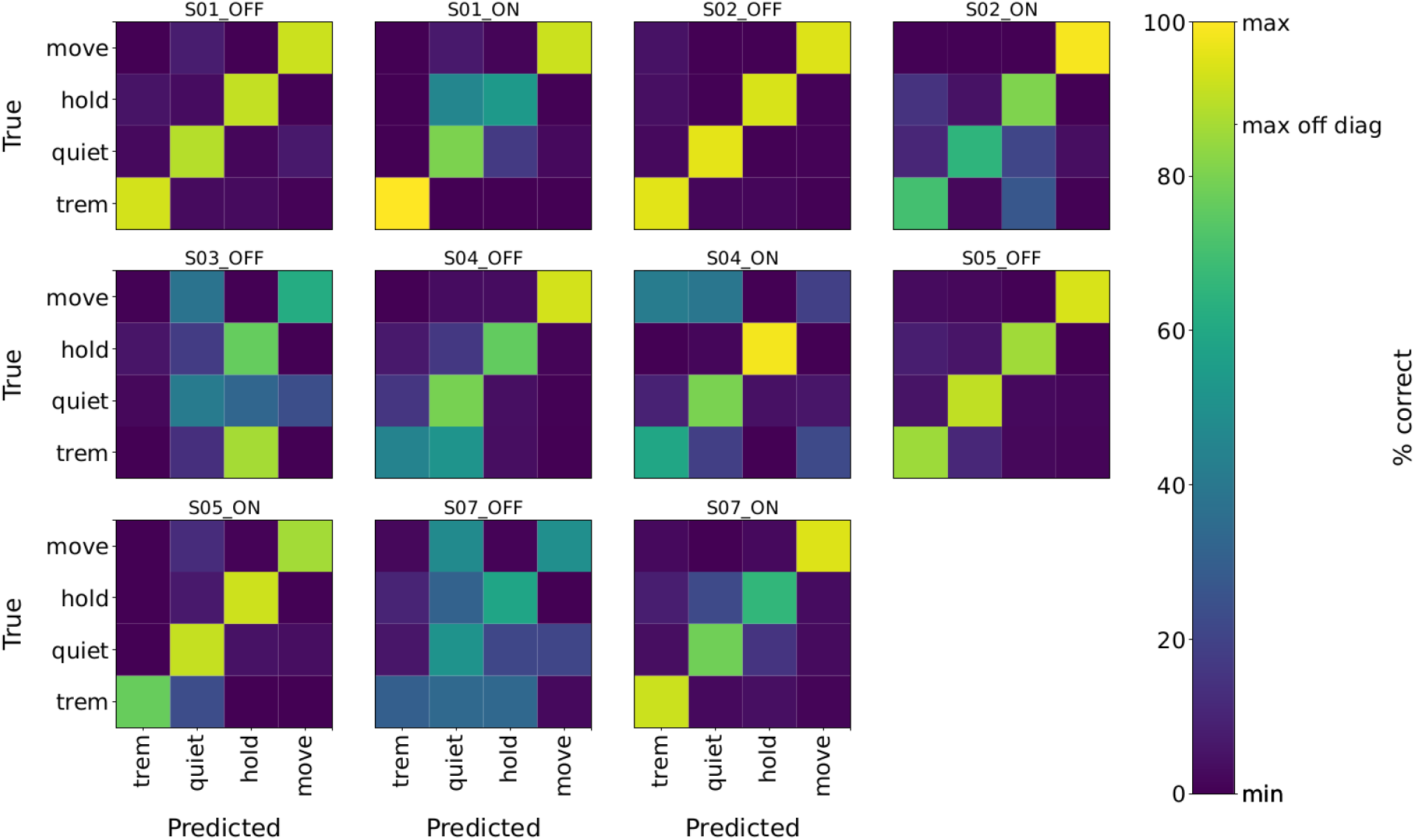
Distinguishing movements based on subthalamic and cortical activity. Confusion matrices for Med On and Med Off per subject, averaged across cross-validation folds. The color bar shows the overall maximum, the maximum of all off-diagonal entries and the minimum of all diagonal entries.

When the model was trained across rather than within subjects, performance of all-to-all classification using STN LFP and MEG data dropped considerably, to 24% in medication OFF and to 28% in medication ON condition (compare Tab. 3), suggesting that movement-specific neural signatures vary across patients.

### STN-cortex two-feature models

While it would be difficult for an adaptive DBS system to monitor the entire cortex, it is feasible to monitor the STN and individual cortical areas (Opri et al., 2020; Gilron et al., 2021). Thus, we investigated the performance gain achieved by adding the activity of a single cortical area to STN activity as a second feature. Fig. 4 shows a brain map of the performance gain. In most cases, sensorimotor cortical areas added the most information.

**Figure 4:**
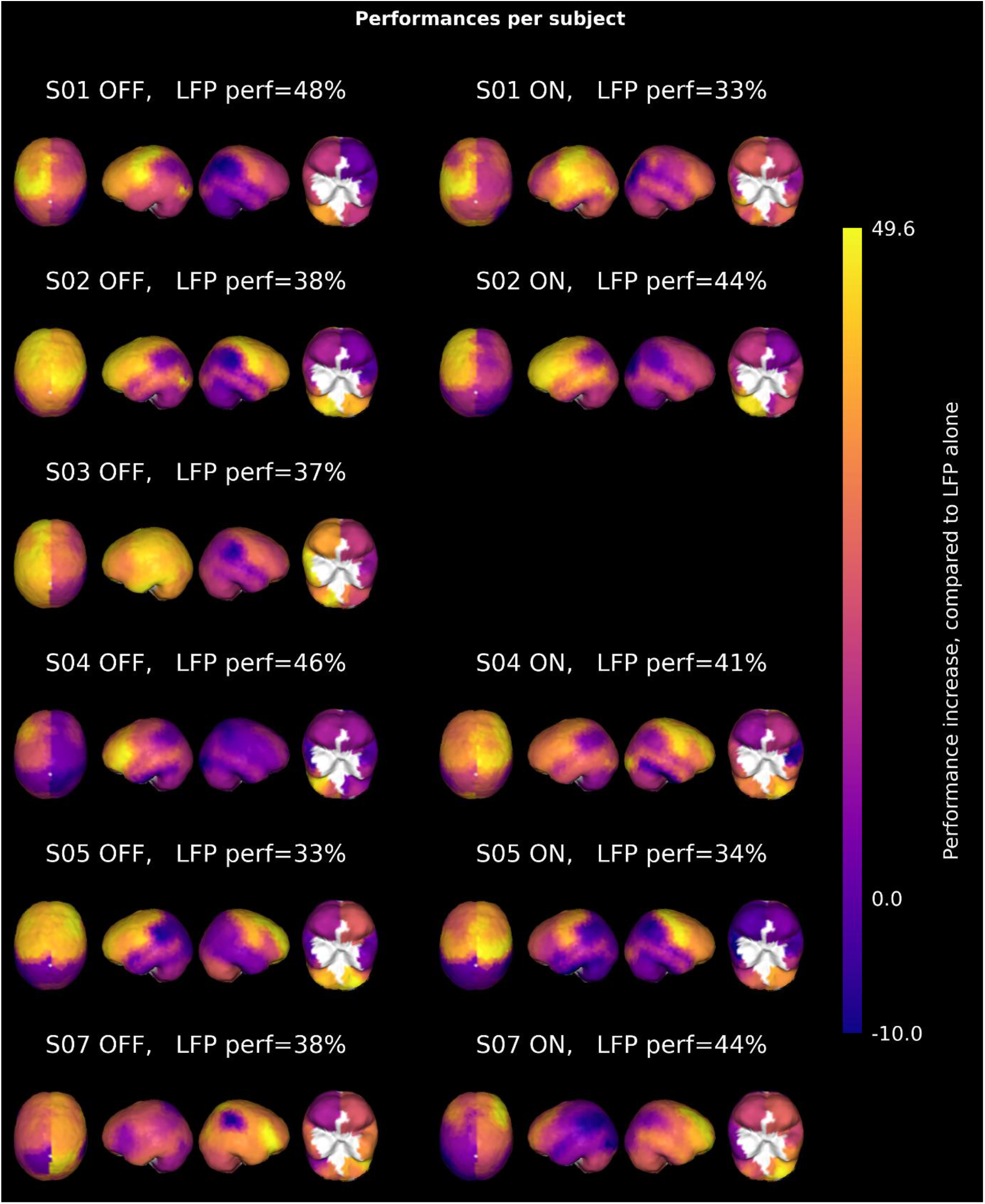
Gain of using cortical activity in addition subthalamic nucleus activity. The balanced accuracy achieved with subthalamic activity alone is provided in the headings (“LFP perf”). Colors code the improvement (in %) obtained by adding the activity of an individual cortical brain area as a second feature.

Adding a single cortical area was usually enough to cover much of the total gain achievable by adding all cortical areas (Fig. 5). Adding all remaining cortical areas to the STN-cortex two-feature models as additional features led to an average performance increase of only 10% (std 11%) compared to the two-feature model.

**Figure 5:**
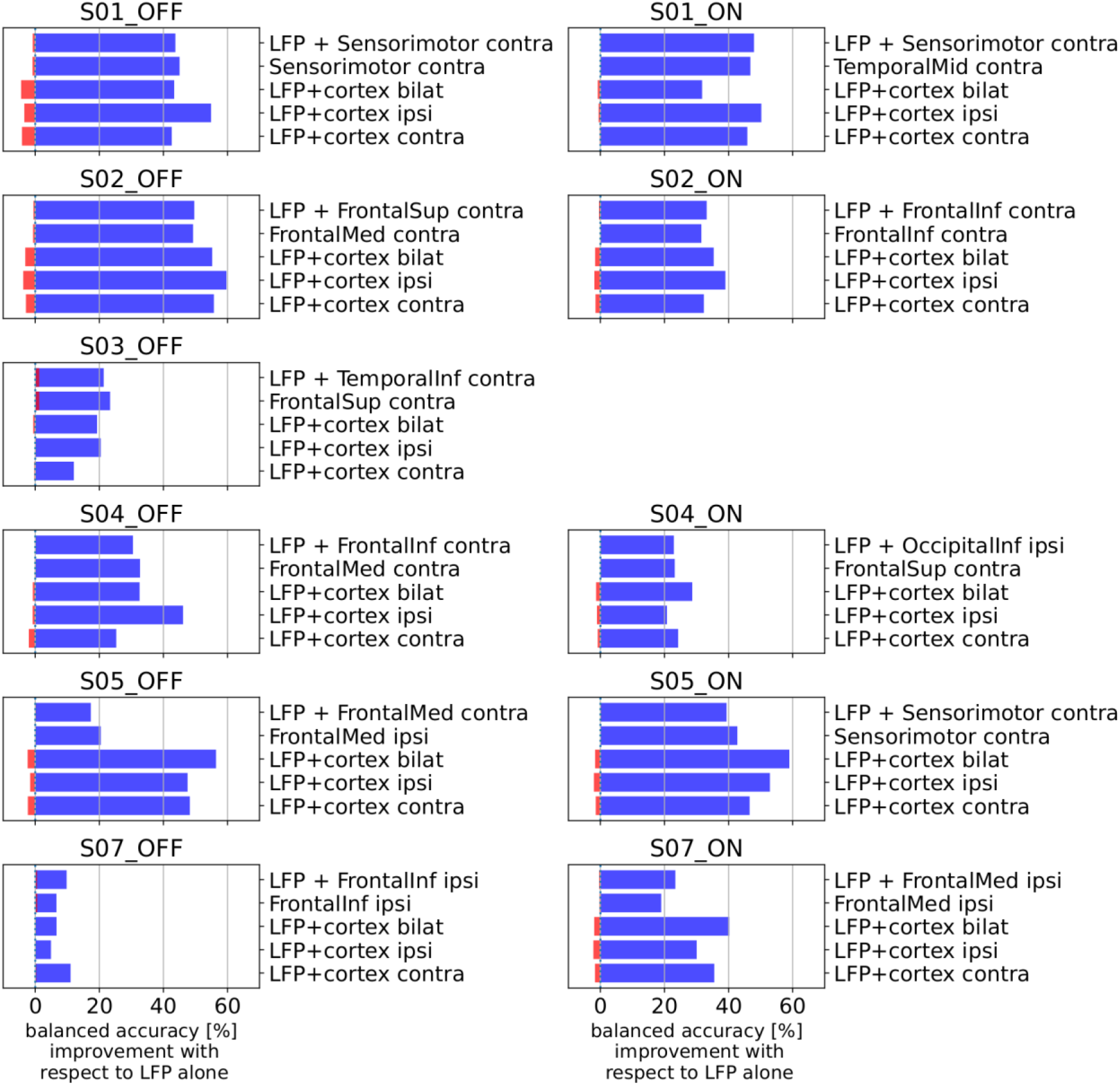
Most informative cortical areas and lateralization of information. Bar height indicates the performance difference with respect to the STN-only case, with original (blue) and shuffled class labels (red). Uppermost bar: best combination of one STN feature and one cortical feature. Second bar: best single cortical feature (STN not used). The lower three bars illustrate the gain of adding all cortical areas (both hemispheres), only the ipsilateral areas (with respect to the moving hand) and only the contralateral areas, respectively.

The spatial distribution of information was not particularly lateralized. The single most informative cortical area was not always on the contralateral side with respect to movement, and adding all contralateral areas to STN activity did not consistently yield higher accuracy than adding all ipsilateral areas (Fig. 5).

## Discussion

In this paper, we demonstrate the feasibility of distinguishing tremor from voluntary movements of the same hand. We found that LFP recordings from the STN are not sufficient for this distinction. It becomes possible, however, when considering additional cortical signals.

### Previous studies

Distinguishing different movements in neural recordings has been the aim of several brain-computer interface studies, typically addressing cortical activity, as measured by EEG (Xu et al., 2021) or electrocorticography (ECoG; Volkova et al., 2019). More recently, the possibility of recording subcortical activity in the context of DBS has facilitated classification studies in basal ganglia and thalamus, most of which deal with movement detection, i.e. the distinction between movement and rest. Loukas and Brown were the first to decode the onset of voluntary movements from subthalamic activity (Loukas and Brown, 2004). Several other studies aimed for tremor detection, due to its relevance for adaptive DBS (Bakstein et al., 2012; Camara et al., 2015; Hirschmann et al., 2017; Shah et al., 2018; Yao et al., 2020). Our group has previously applied Hidden Markov Modelling to STN LFPs, using a superset of the data analyzed here (Hirschmann et al., 2017). Spontaneously occurring PD rest tremor was detected with an average sensitivity of 70% and a specificity of 89%. Yao et al. achieved a sensitivity of 89% and a specificity of 50%, using XGBoost and Kalman filtering of selected input features (Yao et al., 2020). The same group later demonstrated the feasibility of XGBoost tremor detection on chip, facilitating its integration into small, battery-powered neurostimulators (Zhu et al., 2020).Tremor detection has also been demonstrated for thalamic LFPs recorded in ET patients (Tan et al., 2019), including the feasibility of conditioning DBS on these signals under laboratory conditions (He et al., 2021).

In this study, we obtained comparable tremor detection performance for the multifeature case (STN+cortex). Importantly, however, this work goes a step further by distinguishing between tremor, tremor-free rest, self-paced fist-clenching and static forearm extension.

### Clinical implications

Distinguishing between tremor and voluntary hand movements of the same limb based on brain activity is a challenging task, which has both clinical and basic science implications. From a clinical perspective, mastering this task would allow for better adaptation of DBS to the current situation, as it would reduce unnecessary stimulation occurring when an adaptive DBS system confuses voluntary movement, or dyskinesia, with tremor. Unnecessary stimulation is not only a waste of energy, but might contribute to maladaptive changes in response to DBS (Reich et al., 2016). In addition, an unnecessary sudden onset of stimulation or a sudden amplitude increase may interfere with fine motor control, as both may elicit transient side-effects such as paresthesia.

### Neuroscientific implications

From a basic science perspective, it is warranted to understand the neurophysiological differences between voluntary and non-voluntary movements such as tremor. Tremor and voluntary movement share a common oscillatory signature, characterized by beta desynchronization and gamma synchronization (Pfurtscheller et al., 2003; Wang et al., 2005; Beudel et al., 2015; Qasim et al., 2016). While this pattern is very robust, it signals the presence of any movement rather than what movement is being executed, nor whether the movement is voluntary or not. The latter question is highly interesting for general neuroscience and has been addressed by contrasting tremor and mimicked tremor. Although some differences emerged, there were generally more similarities than differences (Muthuraman et al., 2012, 2018). Hence, there is currently no consensus on the neural mechanisms underlying involuntary movement and hence no compelling explanation of why tremor cannot be stopped at will.

Even without a neural correlate of volition, it may be possible to distinguish tremor from other movements based on brain signals. A smart DBS system could simply try to detect oscillatory movements of a certain frequency and assume tremor, as rhythmic movements of this kind are rarely voluntary. This appears to be a feasible approach, given the presumably unique neural correlates of such movements, such as an increase of power at individual tremor frequency (Timmermann et al., 2003; Pollok et al., 2004; Hirschmann et al., 2013a). The characteristic kinematics of tremor and its reflection in brain activity is most likely the reason why were able to distinguish it from other movements here. Note, however, that our approach was based on Hjorth activity, which is not frequency-resolved. Thus, knowing the individual tremor frequency is not necessary for tremor discrimination.

### Movement-related information in STN and cortex

One of the key findings of this paper is that cortical signals can dramatically increase tremor discrimination. This observation tallies with recent studies in PD patients decoding grip force from STN and primary motor cortex, the latter recorded through intraoperative ECoG (Merk et al., 2022; Peterson et al., 2022). Both studies achieved much better predictions when using cortical signals, as compared to subthalamic signals. The authors restrained from drawing conclusions about the information content of cortex versus STN because ECoG signals have a better signal-to-noise ration than recordings made with a DBS electrode, which might explain the difference in performance. In our case, however, the invasive deep brain recording is generally considered to have better signal-to-noise ratio than the noninvasive MEG, suggesting that the STN might not be the optimal brain area for inferring the kind of movement being performed. In fact, a number of studies indicate that the basal ganglia control movement vigor, i.e. the strength and velocity of a movement, rather than being concerned with movement coordination (Turner and Desmurget, 2010; Dudman and Krakauer, 2016; Yttri and Dudman, 2016; Lofredi et al., 2018). While this is in line with the poor decoding of movement type based on STN activity alone, as observed here and in a different analysis of the same data (Hirschmann et al., 2017), it is challenged by recent findings of Golshan and colleagues, who managed to decode different voluntary movements from STN LFPs with accuracies of up to 90% (Golshan et al., 2018, 2020). These studies did not investigate tremor, however, and the approach requires knowing when a movement is initiated. This information, of course, would not be available when applying adaptive DBS in practice.

So as of now, it seems that any smart DBS system that needs know to more about a patient’s motor state than whether or not the patient is at rest would likely profit substantially from external information, such as peripherals (Malekmohammadi et al., 2016b; Cagnan et al., 2017; Cernera et al., 2021) or electrodes in additional brain areas. If so, how many and which brain areas should be monitored? Our results suggest that a few areas suffice for distinguishing tremor from voluntary movement and between different voluntary movements. Unsurprisingly, primary and premotor areas were found to be most useful, although other areas contributed information as well. Thus, a system monitoring both STN and motor cortex appears promising for real-time motor state discrimination in PD.

In fact, such systems have already been tested both in-clinic and at home. Using an implantable DBS system capable of recording neural activity and a combination of deep brain electrodes and cortical strip electrodes placed on primary motor cortex, Gilron et al. found that conditioning STN DBS on cortical gamma oscillations increased the time spent in the Med On/Stim On state without experiencing dyskinesia (Gilron et al., 2021). This is a crucial finding, since reducing stimulation-induced side effects is a key promise of adaptive DBS. Using a similar approach, Opri et al. demonstrated that conditioning thalamic DBS on motorcortical low-frequency activity is as effective for suppressing essential tremor as standard DBS, despite delivering less energy (Opri et al., 2020).

### Generalizability of movement decoding

To be useful in practice, a decoding approach needs to work equally well in different situations, e.g. before and after intake of anti-Parkinsonian medication. In agreement with (Golshan et al., 2020), we obtained similar decoding performance in medication On and Off, suggesting robustness to changes in the level of dopamine. This needs to be tested further, however, in light of recent findings linking decoding performance to motor impairment (Merk et al., 2022; Peterson et al., 2022), which is closely linked to dopamine availability.

Ideally, a decoder would not only generalize to different dopamine levels but to different patients as well. As in almost all previous works on this topic (but see Hirschmann et al., 2022), our approach required within-patient training, i.e. a decoder working in one patient does not necessarily work in another. Tuning the decoder to a new patient requires individual, electrophysiological data from the DBS target and cortex. While this seemed infeasible a few years ago, the advent of new DBS systems capable of recording brain signals in addition to delivering stimulation has made this scenario more realistic (Jimenez-Shahed, 2021). Note, however, that a general decoder would still be preferable from both a practical point of view (no tuning necessary) and a basic science perspective because only general decoders allow for conclusions on the general mechanism of tremor.

### Limitations

One obvious limitation of this work is that we did not detect tremor in real-time nor condition DBS on tremor. In addition, the performance achieved here is not enough for clinical application, although much better than chance, as evidenced by the decline caused by shuffling class labels randomly. Further, the cortical signal was recorded with MEG rather than ECoG. While MEG has much worse spatial resolution, it offers whole-brain coverage. This allows us to exclude movement artifacts as the true source of information because artifacts have a different topography than the information maps provided here (Kandemir et al., 2020). While we found most information in sensorimotor regions, DBS hardware-related artifacts are centered on the adapter for the externalized lead (right parietal and right temporal regions) whereas movement-related MEG artifacts are strongest close to the moving limb.

Finally, we included only men in this re-analysis because they happened to fulfill the inclusion criteria accidentally (spontaneous waxing and waning of tremor in the limb used in the motor task and presence of both hold and grasp movements in the recordings). Hence, we cannot be sure that our findings are transferable to females. Yet, we see no reason for assuming gender differences.

## Conclusions

PD rest tremor can be distinguished from voluntary hand movements when considering subthalamic and cortical signals. This distinction is possible regardless of the dopamine level.

## Funding

This work was supported by Brunhilde Moll Stiftung. Dmitrii Todorov was supported by European Commissions’ MSCA-IF-2017 Grant 793834.

## Acknowledgements

Dmitrii Todorov acknowledges helpful advice about data management and processing from Alexandra Steina and Marius Kroesche. Computations were largely made possible thanks to Human Brain Project ICEI computing resources (project icei-hpb-2020-0012).

## Declaration of interests

None

## Notes

### Competing Interest Statement

The authors have declared no competing interest.

## References

Bakstein E, Burgess J, Warwick K, Ruiz V, Aziz T, Stein J (2012) Parkinsonian tremor identification with multiple local field potential feature classification. J Neurosci Methods 209:320–330.

Beudel M, Little S, Pogosyan A, Ashkan K, Foltynie T, Limousin P, Zrinzo L, Hariz M, Bogdanovic M, Cheeran B, Green AL, Aziz TZ, Thevathasan W, Brown P (2015) Tremor Reduction by Deep Brain Stimulation Is Associated With Gamma Power Suppression in Parkinson’s Disease. Neuromodulation 18:349–354.

Cagnan H, Pedrosa D, Little S, Pogosyan A, Cheeran B, Aziz T, Green A, Fitzgerald J, Foltynie T, Limousin P, Zrinzo L, Hariz M, Friston KJ, Denison T, Brown P (2017) Stimulating at the right time: phase-specific deep brain stimulation. Brain 140:132–145.

Camara C, Isasi P, Warwick K, Ruiz V, Aziz T, Stein J, Bakštein E (2015) Resting tremor classification and detection in Parkinson’s disease patients. Biomed Signal Process Control 16:88–97.

Cernera S, Alcantara JD, Opri E, Cagle JN, Eisinger RS, Boogaart Z, Pramanik L, Kelberman M, Patel B, Foote KD, Okun MS, Gunduz A (2021) Wearable sensor-driven responsive deep brain stimulation for essential tremor. Brain Stimul 14:1434–1443.

Chen T, Guestrin C (2016) XGBoost: A scalable tree boosting system. In: Proceedings of the ACM SIGKDD International Conference on Knowledge Discovery and Data Mining.

Dale J, Schmidt SL, Mitchell K, Turner DA, Grill WM (2022) Evoked potentials generated by deep brain stimulation for Parkinson’s disease. Brain Stimul 15:1040–1047.

Dudman JT, Krakauer JW (2016) The basal ganglia: from motor commands to the control of vigor. Curr Opin Neurobiol 37:158–166.

Gilron R et al. (2021) Long-term wireless streaming of neural recordings for circuit discovery and adaptive stimulation in individuals with Parkinson’s disease. Nat Biotechnol.

Golshan HM, Hebb AO, Hanrahan SJ, Nedrud J, Mahoor MH (2018) A hierarchical structure for human behavior classification using STN local field potentials. J Neurosci Methods.

Golshan HM, Hebb AO, Mahoor MH (2020) LFP-Net: A deep learning framework to recognize human behavioral activities using brain STN-LFP signals. J Neurosci Methods 335.

Gramfort, Alexandre, Martin Luessi, Eric Larson, Denis A. Engemann, Daniel Strohmeier, Christian Brodbeck, Lauri Parkkonen, and Matti S. Hämäläinen. 2014. “MNE Software for Processing MEG and EEG Data.”NeuroImage 86 (February): 446–60. https://doi.org/10.1016/j.neuroimage.2013.10.027.

He S, Baig F, Mostofi A, Pogosyan A, Debarros J, Green AL, Aziz TZ, Pereira E, Brown P, Tan H (2021) Closed-Loop Deep Brain Stimulation for Essential Tremor Based on Thalamic Local Field Potentials. Mov Disord 36.

Hirschmann J, Butz M, Hartmann CJ, Hoogenboom N, Özkurt TE, Vesper J, Wojtecki L, Schnitzler A (2016) Parkinsonian Rest Tremor Is Associated With Modulations of Subthalamic High-Frequency Oscillations. Mov Disord 31:1551–1559.

Hirschmann J, Hartmann CJ, Butz M, Hoogenboom N, Özkurt TE, Elben S, Vesper J, Wojtecki L, Schnitzler A (2013a) A direct relationship between oscillatory subthalamic nucleus-cortex coupling and rest tremor in Parkinson’s disease. Brain 136:3659–3670.

Hirschmann J, Özkurt TE, Butz M, Homburger M, Elben S, Hartmann CJ, Vesper J, Wojtecki L, Schnitzler A (2013b) Differential modulation of STN-cortical and cortico-muscular coherence by movement and levodopa in Parkinson’s disease. Neuroimage 68:203– 213.

Hirschmann J, Schoffelen JM, Schnitzler A, van Gerven MAJ (2017) Parkinsonian rest tremor can be detected accurately based on neuronal oscillations recorded from the subthalamic nucleus. Clin Neurophysiol 128:2029–2036.

Hirschmann J, Steina A, Vesper J, Florin E, Schnitzler A (2022) Neuronal oscillations predict deep brain stimulation outcome in Parkinson’s disease. Brain Stimul 15:792–802.

Jimenez-Shahed J (2021) Device profile of the percept PC deep brain stimulation system for the treatment of Parkinson’s disease and related disorders. Expert Rev Med Devices 18.

Kandemir AL, Litvak V, Florin E (2020) The comparative performance of DBS artefact rejection methods for MEG recordings. Neuroimage 219.

Kelleher JD, Namee B Mac, D’Arcy A (2015) Fundamentals of Machine Learning for Predictive Data Analytics.

Koeglsperger T, Palleis C, Hell F, Mehrkens JH, Bötzel K (2019) Deep brain stimulation programming for movement disorders: Current concepts and evidence-based strategies. Front Neurol 10.

Krauss JK, Lipsman N, Aziz T, Boutet A, Brown P, Chang JW, Davidson B, Grill WM, Hariz MI, Horn A, Schulder M, Mammis A, Tass PA, Volkmann J, Lozano AM (2021) Technology of deep brain stimulation: current status and future directions. Nat Rev Neurol 17.

Lemaître G, Nogueira F, Aridas CK (2017) Imbalanced-learn: A python toolbox to tackle the curse of imbalanced datasets in machine learning. J Mach Learn Res 18.

Little S, Beudel M, Zrinzo L, Foltynie T, Limousin P, Hariz M, Neal S, Cheeran B, Cagnan H, Gratwicke J, Aziz TZ, Pogosyan A, Brown P (2016) Bilateral adaptive deep brain stimulation is effective in Parkinson’s disease. J Neurol Neurosurg Psychiatry 87:717– 721.

Little S, Pogosyan A, Neal S, Zavala B, Zrinzo L, Hariz M, Foltynie T, Limousin P, Ashkan K, FitzGerald J, Green AL, Aziz TZ, Brown P (2013) Adaptive deep brain stimulation in advanced Parkinson disease. Ann Neurol 74:449–457.

Lofredi R, Neumann WJ, Bock A, Horn A, Huebl J, Siegert S, Schneider GH, Krauss JK, Kuühn AA (2018) Dopamine-dependent scaling of subthalamic gamma bursts with movement velocity in patients with Parkinson’s disease. Elife 7.

Loukas C, Brown P (2004) Online prediction of self-paced hand-movements from subthalamic activity using neural networks in Parkinson’s disease. J Neurosci Methods 137:193–205.

Malekmohammadi M, Herron J, Velisar A, Blumenfeld Z, Trager MH, Chizeck HJ, Brontë-Stewart H (2016a) Kinematic Adaptive Deep Brain Stimulation for Resting Tremor in Parkinson’s Disease. Mov Disord 31:426–428.

Malekmohammadi M, Herron J, Velisar A, Blumenfeld Z, Trager MH, Chizeck HJ, Brontë-Stewart H (2016b) Kinematic Adaptive Deep Brain Stimulation for Resting Tremor in Parkinson’s Disease. Mov Disord 31:426–428.

Marceglia S, Guidetti M, Harmsen IE, Loh A, Meoni S, Foffani G, Lozano AM, Volkmann J, Moro E, Priori A (2021) Deep brain stimulation: Is it time to change gears by closing the loop? J Neural Eng 18.

Meidahl AC, Tinkhauser G, Herz DM, Cagnan H, Debarros J, Brown P (2017) Adaptive Deep Brain Stimulation for Movement Disorders: The Long Road to Clinical Therapy. Mov Disord 32.

Merk T, Peterson V, Lipski WJ, Blankertz B, Turner RS, Li N, Horn A, Richardson RM, Neumann WJ (2022) Electrocorticography is superior to subthalamic local field potentials for movement decoding in Parkinson’s disease. Elife 11:2021.04.24.441207.

Milosevic L, Kalia SK, Hodaie M, Lozano AM, Popovic MR, Hutchison WD (2018) Physiological mechanisms of thalamic ventral intermediate nucleus stimulation for tremor suppression. Brain 141:2142–2155.

Muthuraman M, Heute U, Arning K, Anwar AR, Elble R, Deuschl G, Raethjen J (2012) Oscillating central motor networks in pathological tremors and voluntary movements. What makes the difference? Neuroimage 60:1331–1339.

Muthuraman M, Raethjen J, Koirala N, Anwar AR, Mideksa KG, Elble R, Groppa S, Deuschl G (2018) Cerebello-cortical network fingerprints differ between essential, Parkinson’s and mimicked tremors. Brain 141:1770–1781.

Neumann WJ, Turner RS, Blankertz B, Mitchell T, Kühn AA, Richardson RM (2019) Toward Electrophysiology-Based Intelligent Adaptive Deep Brain Stimulation for Movement Disorders. Neurotherapeutics 16.

Nolte G (2003) The magnetic lead field theorem in the quasi-static approximation and its use for magnetoencephalography forward calculation in realistic volume conductors. Phys Med Biol 48:3637–3652.

Oostenveld R, Fries P, Maris E, Schoffelen J-M (2011) FieldTrip: Open Source Software for Advanced Analysis of MEG, EEG, and Invasive Electrophysiological Data. Comput Intell Neurosci 2011:1–9.

Opri E, Cernera S, Molina R, Eisinger RS, Cagle JN, Almeida L, Denison T, Okun MS, Foote KD, Gunduz A (2020) Chronic embedded cortico-thalamic closed-loop deep brain stimulation for the treatment of essential tremor. Sci Transl Med 12:eaay7680.

Peterson V, Merk T, Bush A, Nikulin V, Kühn AA, Neumann W-J, Richardson RM (2022) Movement decoding using spatio-spectral features of cortical and subcortical local field potentials. Exp Neurol 359:114261.

Pfurtscheller G, Graimann B, Huggins JE, Levine SP, Schuh LA (2003) Spatiotemporal patterns of beta desynchronization and gamma synchronization in corticographic data during self-paced movement. Clin Neurophysiol 114:1226–1236.

Piña-Fuentes D, Little S, Oterdoom M, Neal S, Pogosyan A, Tijssen MAJ, van Laar T, Brown P, van Dijk JMC, Beudel M (2017) Adaptive DBS in a Parkinson’s patient with chronically implanted DBS: A proof of principle. Mov Disord 32:1253–1254.

Pollok B, Gross J, Dirks M, Timmermann L, Schnitzler A (2004) The cerebral oscillatory network of voluntary tremor. J Physiol 554:871–878.

Qasim SE, de Hemptinne C, Swann N, Miocinovic S, Ostrem JL, Starr PA (2016) Electrocorticography reveals beta desynchronization in the basal ganglia-cortical loop during rest tremor in Parkinson’s disease. Neurobiol Dis 86:177–186.

Reich MM, Brumberg J, Pozzi NG, Marotta G, Roothans J, Åström M, Musacchio T, Lopiano L, Lanotte M, Lehrke R, Buck AK, Volkmann J, Isaias IU (2016) Progressive gait ataxia following deep brain stimulation for essential tremor: adverse effect or lack of efficacy? Brain 139:2948–2956.

Shah SA, Tinkhauser G, Chen CC, Little S, Brown P (2018) Parkinsonian Tremor Detection from Subthalamic Nucleus Local Field Potentials for Closed-Loop Deep Brain Stimulation. Proc Annu Int Conf IEEE Eng Med Biol Soc EMBS 2018-July:2320–2324.

Swann NC, De Hemptinne C, Thompson MC, Miocinovic S, Miller AM, Gilron R, Ostrem JL, Chizeck HJ, Starr PA (2018) Adaptive deep brain stimulation for Parkinson’s disease using motor cortex sensing. J Neural Eng 15.

Tan H, Debarros J, He S, Pogosyan A, Aziz TZ, Huang Y, Wang S, Timmermann L, Visser-Vandewalle V, Pedrosa DJ, Green AL, Brown P (2019) Decoding voluntary movements and postural tremor based on thalamic LFPs as a basis for closed-loop stimulation for essential tremor. Brain Stimul 12:858–867.

Taulu, S, and J Simola. 2006. “Spatiotemporal Signal Space Separation Method for Rejecting Nearby Interference in MEG Measurements.” Physics in Medicine and Biology 51 (7): 1759–68. https://doi.org/10.1088/0031-9155/51/7/008.

Timmermann L, Gross J, Dirks M, Volkmann J, Freund HJ, Schnitzler A (2003) The cerebral oscillatory network of parkinsonian resting tremor. Brain 126:199–212.

Tinkhauser G, Pogosyan A, Little S, Beudel M, Herz DM, Tan H, Brown P (2017) The modulatory effect of adaptive deep brain stimulation on beta bursts in Parkinson’s disease. Brain 140.

Turner RS, Desmurget M (2010) Basal ganglia contributions to motor control: a vigorous tutor. Curr Opin Neurobiol 20:704–716.

Tzourio-Mazoyer N, Landeau B, Papathanassiou D, Crivello F, Etard O, Delcroix N, Mazoyer B, Joliot M (2002) Automated anatomical labeling of activations in SPM using a macroscopic anatomical parcellation of the MNI MRI single-subject brain. Neuroimage 15.

Volkova K, Lebedev MA, Kaplan A, Ossadtchi A (2019) Decoding Movement From Electrocorticographic Activity: A Review. Front Neuroinform 13.

Wang SY, Aziz TZ, Stein JF, Liu X (2005) Time-frequency analysis of transient neuromuscular events: Dynamic changes in activity of the subthalamic nucleus and forearm muscles related to the intermittent resting tremor. J Neurosci Methods 145:151– 158.

Xu L, Xu M, Jung TP, Ming D (2021) Review of brain encoding and decoding mechanisms for EEG-based brain–computer interface. Cogn Neurodyn 15.

Yao L, Brown P, Shoaran M (2020) Improved detection of Parkinsonian resting tremor with feature engineering and Kalman filtering. Clin Neurophysiol 131:274–284.

Yttri EA, Dudman JT (2016) Opponent and bidirectional control of movement velocity in the basal ganglia. Nat 2016 5337603 533:402–406.

Zarzycki MZ, Domitrz I (2020) Stimulation-induced side effects after deep brain stimulation-A systematic review. Acta Neuropsychiatr.

Zhu B, Farivar M, Shoaran M (2020) ResOT: Resource-Efficient Oblique Trees for Neural Signal Classification. IEEE Trans Biomed Circuits Syst 14.

